# FastSpar: Rapid and scalable correlation estimation for compositional data

**DOI:** 10.1101/272583

**Authors:** Stephen C. Watts, Scott C. Ritchie, Michael Inouye, Kathryn E. Holt

**Author notes:** Correspondence to (SCW -).

## Abstract

A common goal of microbiome studies is the elucidation of community composition and member interactions using counts of taxonomic units extracted from sequence data. Inference of interaction networks from sparse and compositional data requires specialised statistical approaches. A popular solution is SparCC, however its performance limits the calculation of interaction networks for very high-dimensional datasets. Here we introduce FastSpar, an efficient and parallelisable implementation of the SparCC algorithm which rapidly infers correlation networks and calculates *p*-values using an unbiased estimator. We further demonstrate that FastSpar reduces network inference wall time by 2-3 orders of magnitude compared to SparCC. FastSpar source code, precompiled binaries, and platform packages are freely available on GitHub: github.com/scwatts/FastSpar

## Introduction

Microbiome analysis, which aims to assay the bacterial communities present in a given sample set, is important in many fields spanning from human health to agriculture and environmental ecology. The current standard for investigating bacterial community composition is to deep sequence the total genomic DNA or the bacterial 16S rRNA gene and analyse the genetic diversity and abundance within each sample. Unique sequences or sequence clusters are taken to represent operational taxonomic units (OTUs) present in the original sample, and the frequencies of these across samples are summarised in the form of an OTU table (Ju and Zhang 2015). In many studies, this data is then exploited to construct correlation networks of OTUs spanning sample sets, which can be used to infer or approximate interactions between taxa (Nakatsu *et al*., 2015, He *et al*., 2017).

The calculation of OTU correlation values is complicated by the sparse and compositional nature of the data. OTU counts are typically normalised by dividing each observation within a sample by the total count for that sample, giving a measure of relative abundance. However this transformation introduces dependencies between normalised sample observations, such that calculating simple correlations from the resulting values is not statistically valid (Aitchison 1982). To perform robust and unbiased statistical analysis of sparse compositional data, it is generally first transformed from the simplex to Euclidean real space.

Returning compositional OTU data back to Euclidean real space can be achieved by taking the log ratio of OTU fractions. Utilising log-ratios restores independence for each OTU and allows components to take on a positive or negative value. Building upon the use of log ratios, Friedman and Alm (2012) articulate an important and robust algorithm, SparCC, to estimate the linear Pearson Correlation between OTUs from variances of log ratios. Given that correlations cannot be calculated directly from log ratio variances, SparCC estimates the correlation statistic by using log ratio variances to approximate the true OTU variance on the assumption that the number of strong correlates is small (Friedman and Alm 2012).

A Python 2 implementation of the SparCC algorithm has been released by the authors with several ancillary scripts for *p*-value estimation. However, the performance of this implementation precludes analysis of large datasets such as those generated from longitudinal studies (Teo *et al*., 2017). Further, the *p*-value estimator used by SparCC has been demonstrated to be biased and overestimate significance (Phipson and Smyth 2010).

Here we present FastSpar, a fast and parallelisable implementation of the SparCC algorithm with an unbiased *p*-value estimator. We demonstrate that FastSpar produces equivalent OTU correlations as SparCC while greatly reducing run time and memory consumption on large data sets. We also show that FastSpar has superior performance to the unpublished re-implementations of SparCC available in the mothur and SpiecEasi packages.

### Implementation

FastSpar is written in C++11, utilising OpenBLAS and LAPACK via the Armadillo library (Xianyi *et al*., 2012; Dongarra *et al*., 1992; Conrad *et al.*, 2016). The GNU Scientific Library (GSL) provides functionality for OTU fraction estimation and threading support is delivered by OpenMP (Dagum *et al.*, 1998). In place of the *p*-value estimator used in SparCC, we utilised an estimator which corrects *p*-value understatement by considering the possibility of recalling repetitious permutations or original data during testing (Phipson and Smyth 2010).

## Results

### Algorithm fidelity

To demonstrate that FastSpar produces equivalent correlations as SparCC, correlation networks were constructed by both programs using random subsets of an OTU table generated from the American Gut Project 16S rRNA sequence data (www.americangut.org), comprising a total of 6,068 OTUs and 7,523 samples. For each OTU pair, the mean correlation values calculated across 20 replicate runs were near identical for FastSpar and SparCC (**Fig. 1** and **2**). The OTU correlations calculated by SparCC and FastSpar are not reproduced exactly as the underlying algorithm begins by non-deterministically estimating OTU fractions. Hence replicate runs of either program on the same input table produce similar but non-identical results (**Fig. 1a-b**). To allow direct comparison of the algorithms, OTU fractions were pre-computed and provided as an additional input to both SparCC and FastSpar (note that the behaviour of the pseudo-random number generators (PRNG) used by FastSpar (GSL) and SparCC (numpy) differ, thus seeding the PRNGs is insufficient to enable direct comparison). When using the same pre-computed OTU fractions as input, FastSpar and SparCC returned identical results (**Fig. 1d**). These comparisons can be reproduced by running the code at github.com/scwatts/fastspar_comparison. The values calculated by the SpiecEasi implementation of SparCC showed similar fidelity to that of FastSpar; however the mothur implementation of SparCC showed much poorer fidelity (**Fig 3**).

**Fig. 1.**
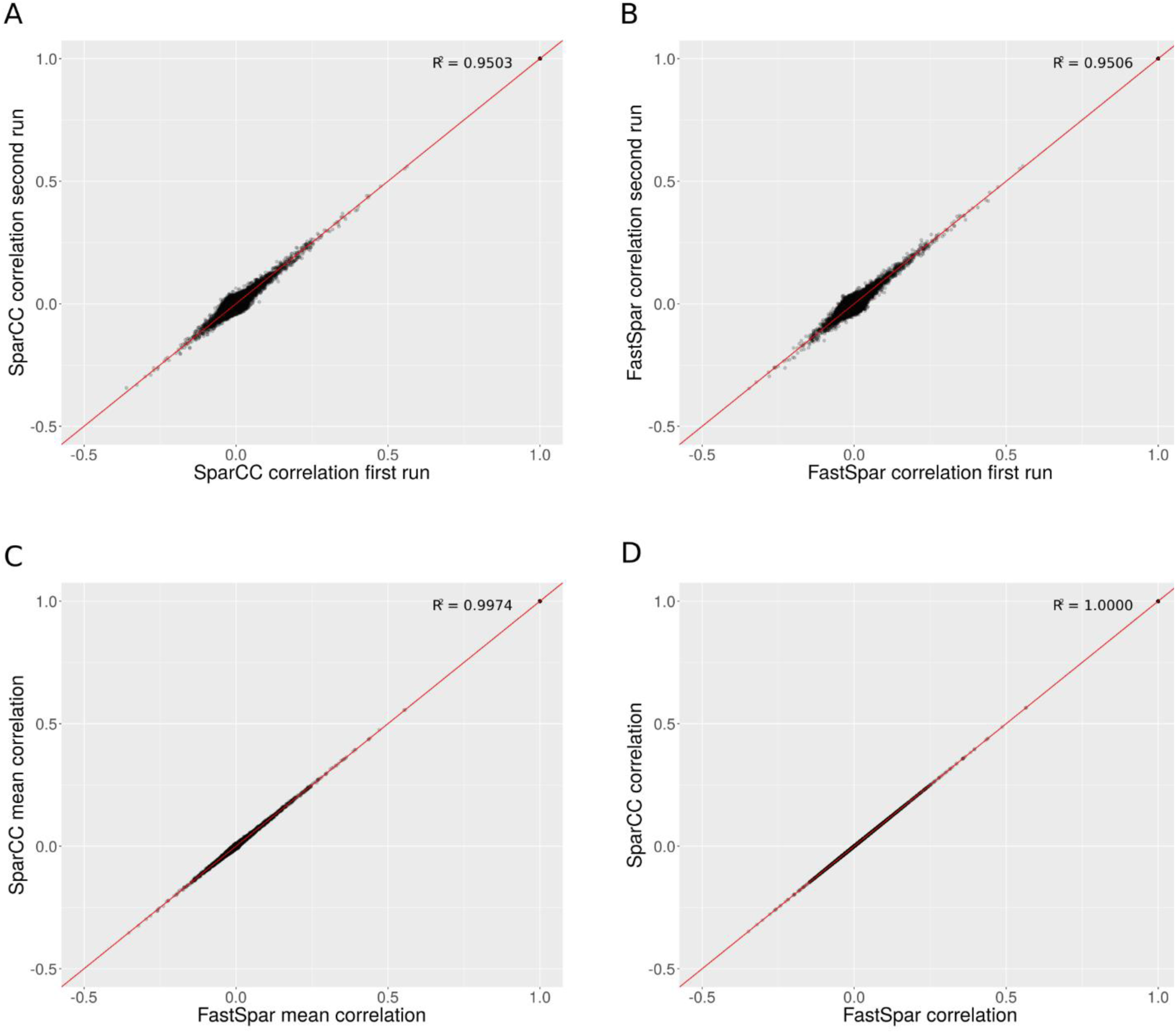
Comparison of OTU correlation values estimated by FastSpar and SparCC. (**A-B**) Pairwise comparison of correlation values estimated across 20 replicate runs using the same implementation (**A**, SparCC or **B**, FastSpar). Note the algorithm is non-deterministic as OTU fractions are drawn from a probability distribution, hence variation of correlation values between replicates runs is observed with either implementation. (**C**) Pairwise comparison of mean estimates across 20 replicate runs, for SparCC vs FastSpar. Note that agreement between the mean estimates of the two implementations is greater than the agreement between replicate runs of the same implementation (panels **A-B**). (**D**) Direct comparison of correlation values generated by SparCC vs FastSpar using the same (i.e. non-random, pre-computed) OTU fractions, showing that FastSpar produces an identical result to SparCC.

**Fig. 2.**
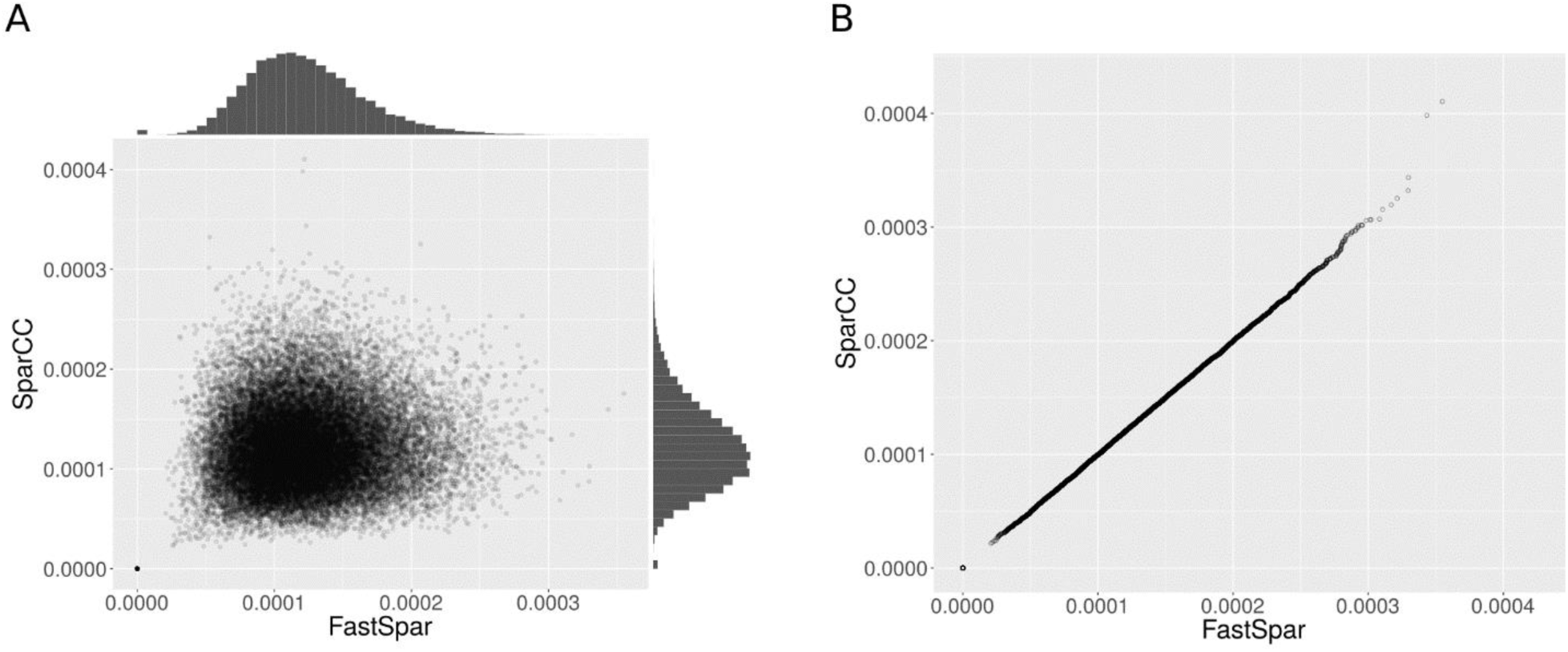
(**A**) Distributions and (**B**) Q-Q plot of pairwise OTU correlation variance in 20 replicate runs of FastSpar and SparCC. OTU correlations were calculated for all pairs of 6,068 OTUs across 7,523 samples.

**Fig. 3.**
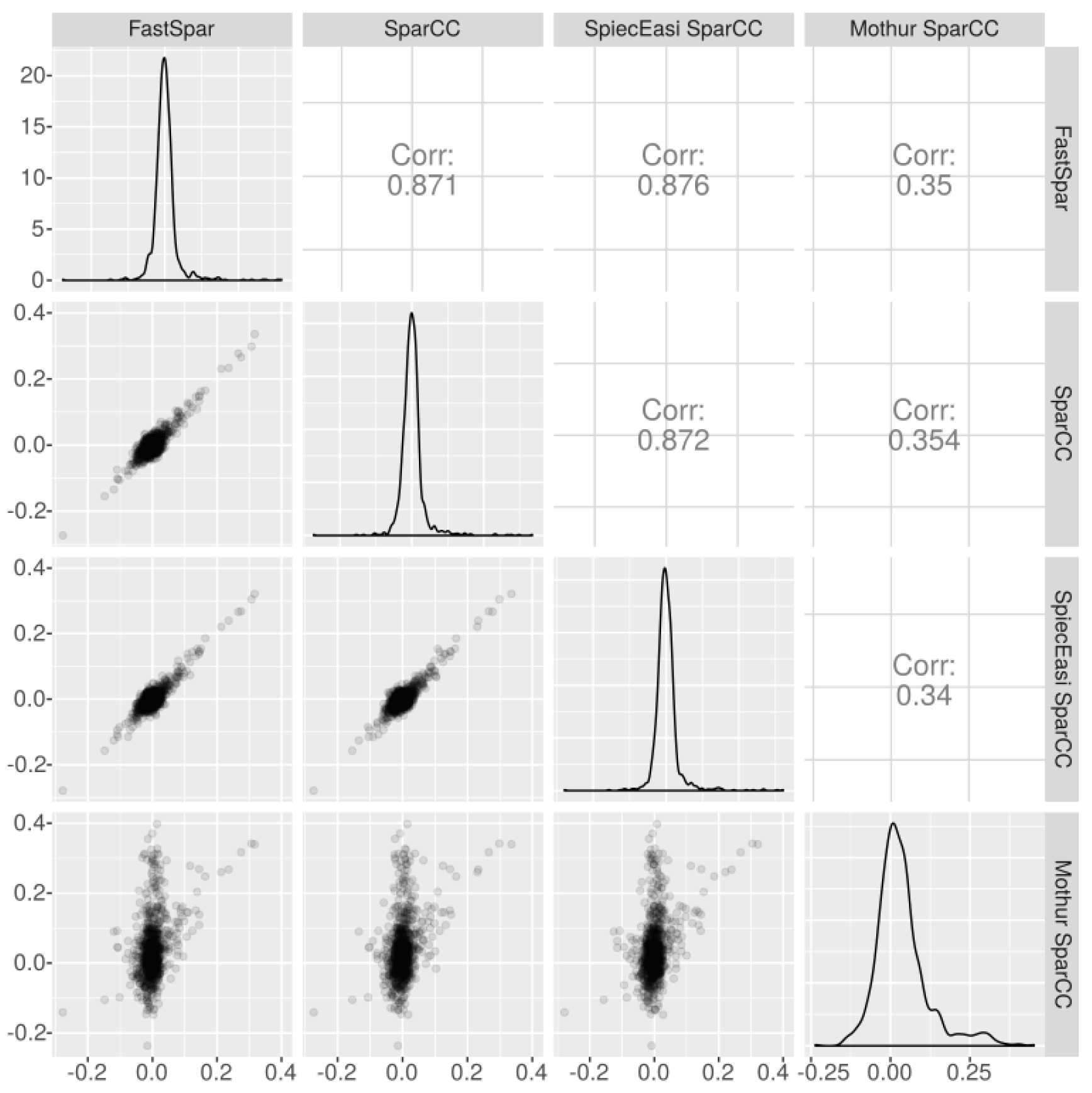
Comparison of OTU correlates calculated by each software package using a random subset of 200 samples and 50 OTUs. Both the FastSpar and SpiecEasi implementation replicate correlations produced by SparCC. OTU correlates calculated by the mothur implementation do not follow the same distribution as those calculated by SparCC.

### Performance profiling

Performance was compared by running FastSpar and SparCC on random subsets of the American Gut Project OTU table (**Fig. 4**). Ten random subsets of each combination of sample sizes (n=250, 500, …, 2500) and OTUs (n=250, 500, …, 2500) were generated, and subjected to analysis using FastSpar (with and without threading) and SparCC. Wall time and memory usage was recorded using GNU time. The analysis was completed in an Ubuntu 17.04 (Zesty Zapus) chroot environment with the required software packages (**Table 1**). Computation was performed with an Intel(R) Xeon(R) CPU E5-2630 @ 2.30GHz CPU and 62 GB RAM. The performance profiling can be reproduced by running the code at github.com/scwatts/fastspar_timing.

**Table 1.** Software packages with version designations used for performance profiling and output comparison.

| Package | Version |
| --- | --- |
| build-essential | 12.1ubuntu2 |
| git | 1:2.7.4-0ubuntu1 |
| libarmadillo-dev | 1:6.500.5+dfsg-1 |
| libgsl-dev | 2.1+dfsg-2 |
| libopenblas-dev | 0.2.18-1ubuntu1 |
| mercurial | 3.7.3-1ubuntu1 |
| python-numpy | 1:1.11.0-1ubuntu1 |
| python-pandas | 0.17.1-3ubuntu2 |
| python3-numpy | 1:1.11.0-1ubuntu1 |
| r-base-core | 3.2.3-4 |
| time | 1.7-25.1 |

**Fig 4.**
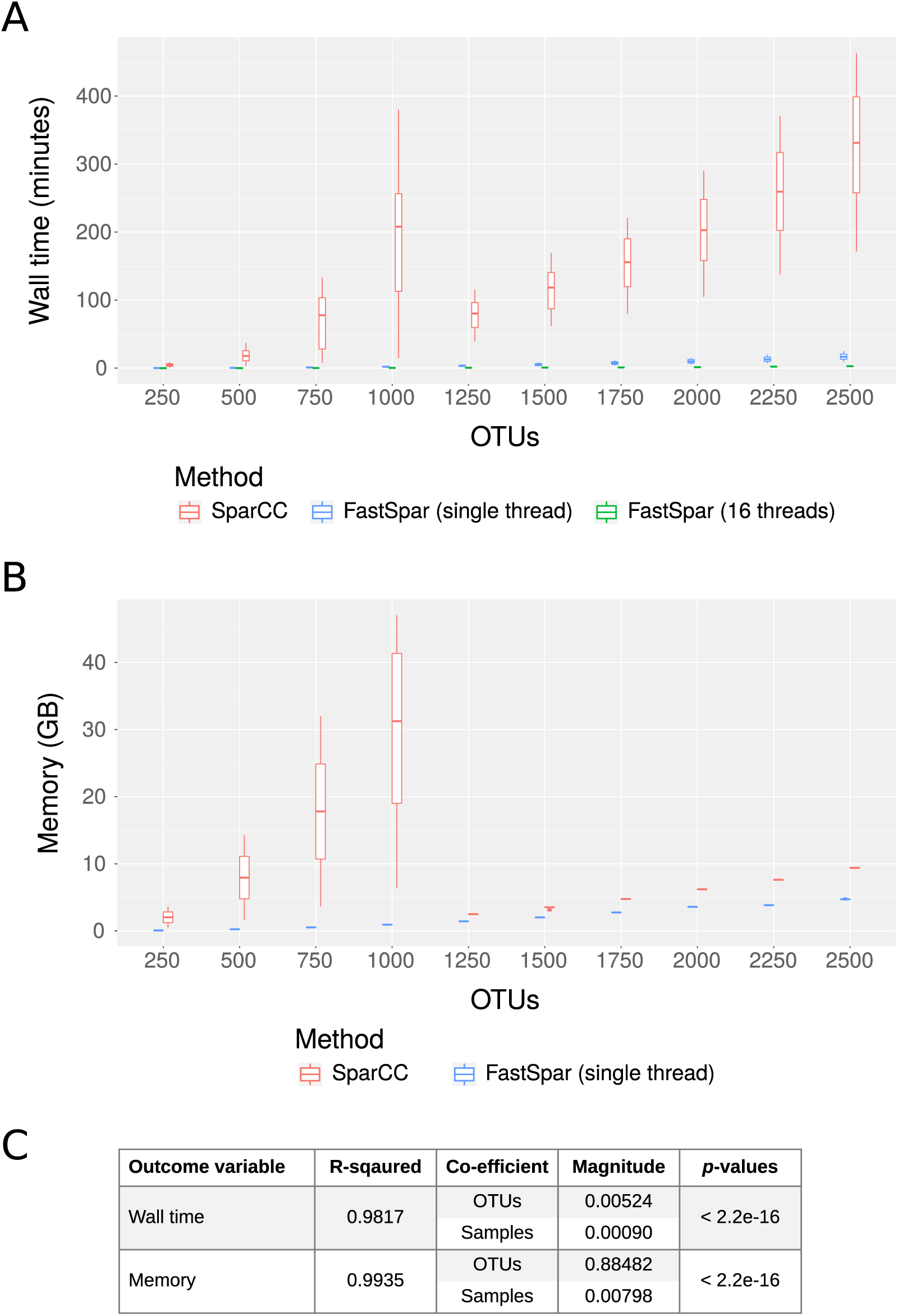
Performance profile of FastSpar and SparCC across random subsets of different sizes, extracted from the American Gut Project OTU table. (**A**) Wall time and (**B**) memory profiles were recorded using GNU time. (**C**) Linear models describing FastSpar (single thread) performance metrics with relation to input data dimensions.

Using 16 threads, FastSpar was up to 821× faster than SparCC, (mean 221× faster; **Fig. 4a**). Using a single thread, FastSpar was up to 118× faster than SparCC (mean 32× faster; **Fig. 4a**). The memory usage of FastSpar was up to 60× less than SparCC (mean 14× less; **Fig. 4b**). Notably the memory performance of SparCC on datasets with 1,000 or more OTUs improves considerably and is due to the conditional use of a more memory efficient calculation for the variation matrix (**Fig. 4b**). This conditional calculation appears to be beneficial for SparCC when analysing datasets with 500 or fewer OTUs but causes a substantial performance degradation for datasets with 500 to 1,000 OTUs (**Fig. 5**). Comparison to the unpublished re-implementations of the SparCC algorithm using an OTU table of consisting of size 500 samples and 1,000 OTUs randomly chosen showed superior performance for FastSpar (**Fig 6**).

**Fig. 5.**
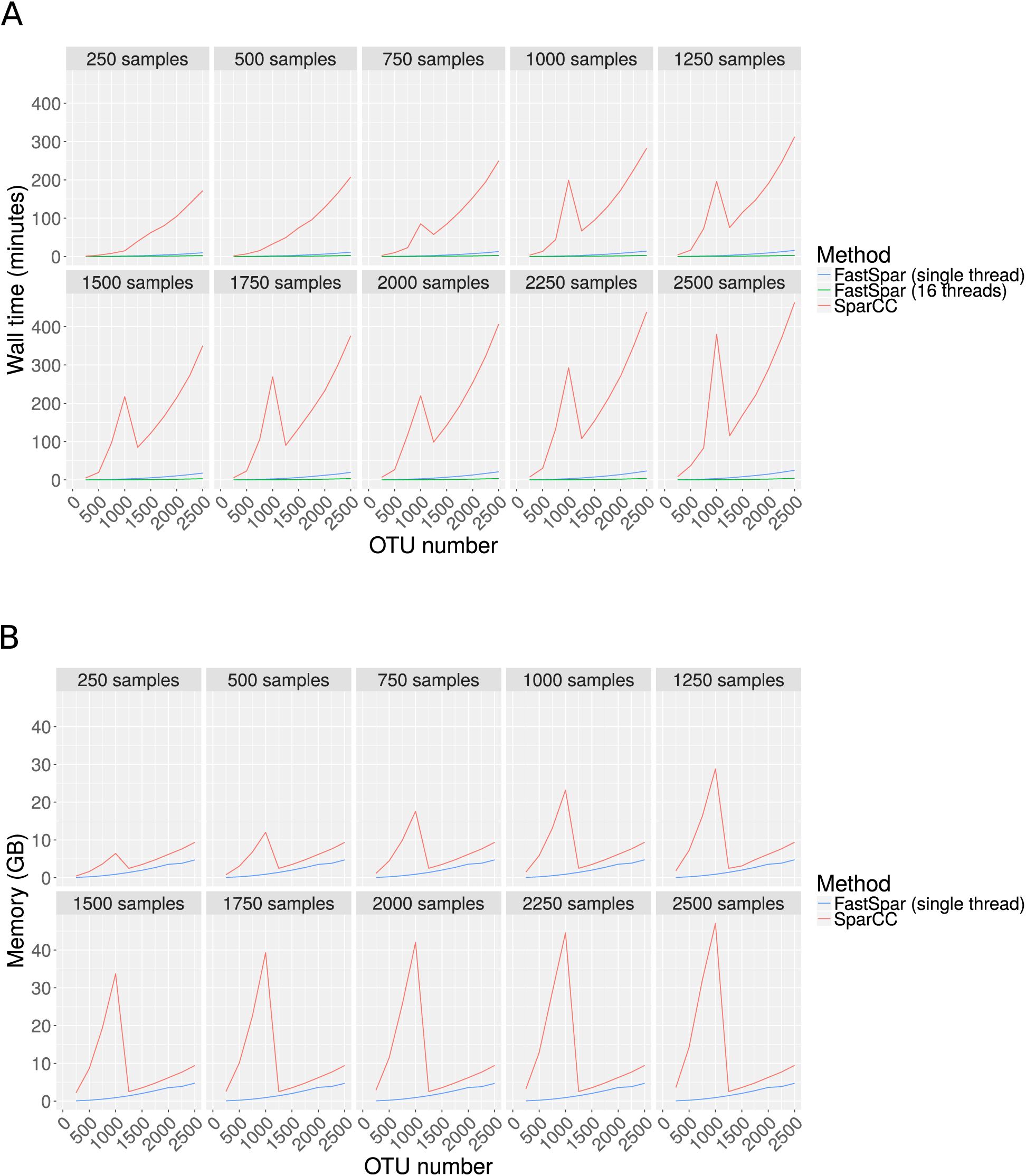
Performance profile of (**A**) FastSpar and (**B**) SparCC for each individual random subset of the American Gut Project OTU table (full table contains 6,068 OTUs and 7,523 samples). Wall time and memory profiles recorded using GNU time.

**Fig. 6.**
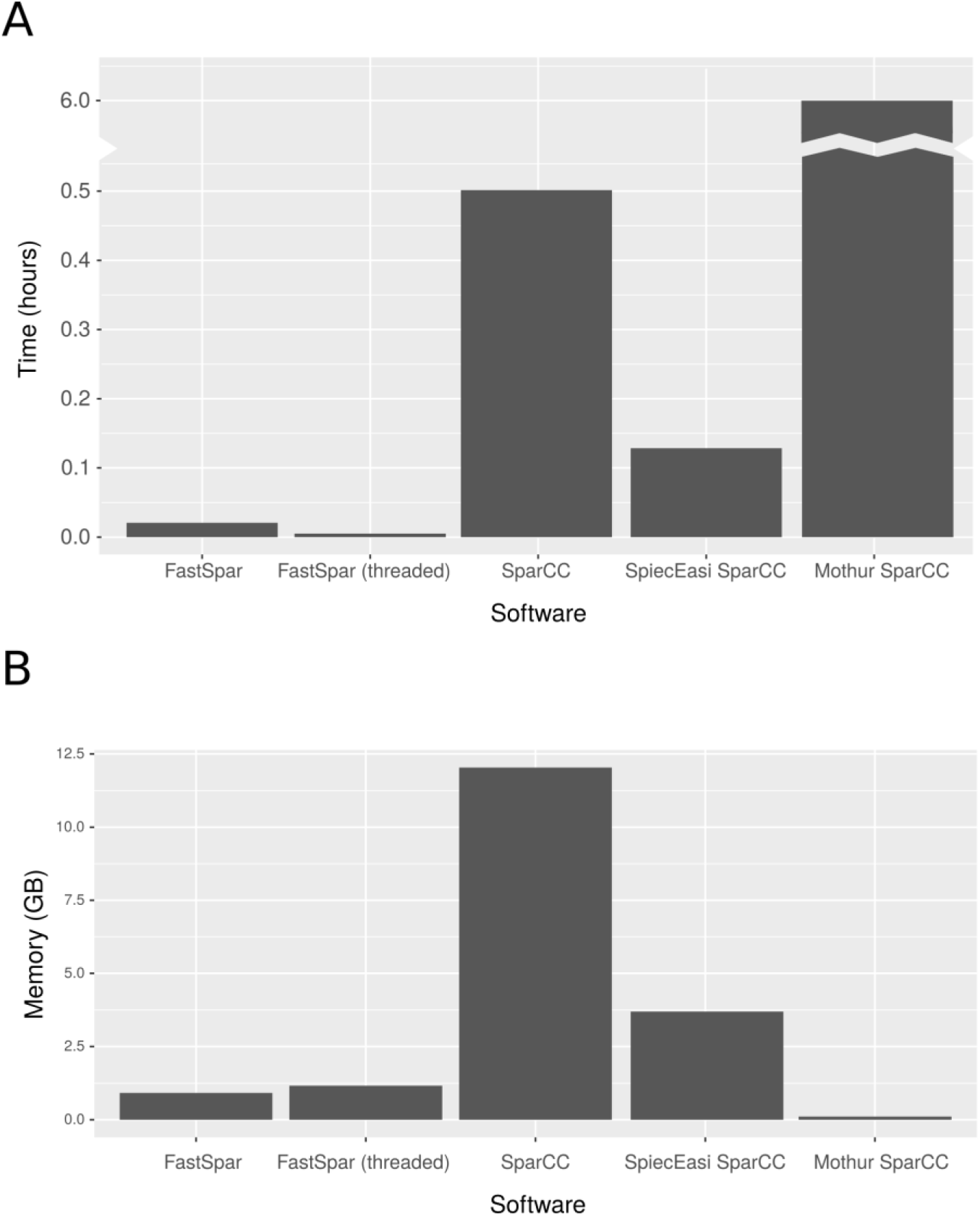
Comparison of **(A)** time and **(B)** memory profiles for existing SparCC algorithm implementations. To perform profiling we used an OTU table composed of 500 samples and 1,000 OTUs randomly selected from the American Gut Project OTU table. FastSpar is demonstrated to have the highest improvement of run time. The SpiecEasi implementation of SparCC shows a moderate performance improvement with regards to both time and memory consumption whereas the mothur implementation did not finish within 6 hours of run time.

As expected, both run time and memory principally scale with OTU number rather than sample number (**Fig. 4c**). For large data sets, it is therefore essential to perform pre-processing of the OTU table in order to reduce the number of OTUs prior to calculating correlations. This can be achieved primarily using two approaches: (1) filtering poorly represented OTUs, or (2) distribution-based clustering such as that used in dbOTU3. The latter approach aims to reunite OTUs derived from sequencing error with the parent OTU by clustering OTUs based on nucleotide edit distance and count distribution (Preheim *et al*. 2013). This has the advantage of retaining count information and thus improving statistical power. To simplify the workflow for large-scale correlation network analyses of microbiome data, FastSpar is packaged with an efficient C++11 implementation of dbOTU3 (github.com/scwatts/dbotu) that has been optimised for analysis of large datasets by applying concurrency design patterns.

FastSpar provides a more robust and efficient method for inferring correlation networks from large microbiome datasets, which was previously intractable yet is likely to become commonplace in modern cohort studies.

## Acknowledgements

This work was supported by the National Health and Medical Research Council of Australia (Project #1062227, Fellowship #1061409 to K.E.H., Fellowship #1061435 to M.I. co-funded by the Australian Heart Foundation) and by the Australian Government Research Training Program (Scholarship to S.W and S.R.).

